# The *Plasmodium* transmission-blocking symbiont, *Microsporidia MB*, is vertically transmitted through *Anopheles arabiensis* germline stem cells

**DOI:** 10.1101/2024.06.14.598976

**Authors:** Thomas O. Onchuru, Edward E. Makhulu, Purity C. Ronnie, Stancy Mandere, Fidel G. Otieno, Joseph Gichuhi, Jeremy K. Herren

## Abstract

*Microsporidia MB* is a promising candidate for developing a symbiont-based strategy for malaria control because it disrupts the capacity of *An. arabiensis* to transmit the *Plasmodium* parasite. The symbiont is predominantly localized in the reproductive organs and is transmitted vertically from mother to offspring and horizontally (sexually) during mating. Due to the contribution of both transmission routes, *Microsporidia MB* has the potential to spread through target vector populations and become established at high prevalence. Stable and efficient vertical transmission of *Microsporidia MB* is important for its sustainable use for malaria control, however, the vertical transmission efficiency of *Microsporidia MB* can vary. In this study, we investigate the mechanistic basis of *Microsporidia MB* vertical transmission in *An. arabiensis*. We show that vertical transmission occurs through the acquisition of *Microsporidia MB* by *Anopheles* cystocyte progenitors following the division of germline stem cells. We also show that *Microsporidia MB* replicates to increase infection intensity in the oocyte of developing eggs when mosquitoes are given a blood meal suggesting that symbiont proliferation in the ovary is coordinated with egg development. The rate of *Microsporidia MB* transmission to developing eggs is on average higher than the recorded (mother to adult offspring) vertical transmission rate. This likely indicates that a significant proportion of *An. arabiensis* offspring lose their *Microsporidia MB* symbionts during development. The stability of germline stem cell infections, coordination of symbiont proliferation, and very high rate of transmission from germline stem cells to developing eggs indicate that *Microsporidia MB* has a highly specialized vertical transmission strategy in *An. arabiensis,* which may explain host specificity.

**Author Summary:** Mosquito vectors of diseases are associated with a broad range of microbes. Some of the microbes significantly affect vector biology including pathogen transmission efficiency. Anopheles mosquitoes, which transmit the malaria parasite, *Plasmodium falciparum,* harbor a native microbe known as *Microsporidia MB.* This microbe interferes with the formation of transmissible stages of the parasite that are transferred to humans by female mosquitoes when taking a blood meal. This phenotype can be exploited to develop a novel strategy for controlling malaria similar to the control of dengue fever using Aedes mosquitoes carrying *Wolbachia* bacteria. Mother-to-offspring transmission of protective microbes is important in sustainable application of microbe-based technologies to control vector-borne diseases because it ensures maintenance of the microbe in target vector populations across many generations. Here, we investigated stability of *Microsporidia MB* infections and efficiency of mother-to-offspring transmission during early stages of egg formation and development. We found that this microbe has a specialized transmission mechanism that involves infecting the germline cells that are important in egg production. We also demonstrated a very high transmission rate (97%) of the *Microsporidia MB* from infected germline cells into daughter cells during cell division. As the germline daughter cells developed into eggs, *Microsporidia MB* established itself in the egg yolk through active replication which only occured after the female mosquitoes had a blood meal. Our study gives insights into an efficient mother-to-offspring transmission route of *Microsporidia MB* that can be utilized sustainably in microbe-based intervention to control malaria.

## Introduction

Vector-borne diseases, and in particular malaria, are of global importance because of the associated economic burden and high mortality rates, especially in developing countries (1–3). Malaria is transmitted by *Anopheles* mosquitoes and is responsible for over half a million deaths annually (3). The primary method for controlling this disease is vector management which includes the use of long-lasting insecticidal nets (LLINs), indoor residual spraying (IRS), and the management of mosquito breeding sites (4). The application of these methods has significantly reduced transmission intensity and subsequently the malaria burden.

However, the progress in malaria control has stalled suggesting that the current malaria control strategies have limitations in driving malaria cases to significantly low levels for successful elimination (3,5). Therefore, the development of novel approaches is urgently needed to sustain the fight against malaria.

There are now several cases where it has been demonstrated that microbes associated with mosquito vectors can significantly alter their ability to efficiently transmit pathogens (Moreira et al., 2009). For instance, the vectorial capacity of the *Aedes aegypti* mosquito, which is responsible for the transmission of medically important viruses such as dengue, chikungunya, and zika, is significantly impaired in the presence of an intracellular endosymbionts, *Wolbachia* (6). *Wolbachia* has now been successfully deployed to control dengue fever transmission (7–9). Similarly, it has been demonstrated that the association of malaria-transmitting mosquitoes and specific microbial symbionts reduces the ability of these vectors to transmit malaria-causing *Plasmodium* parasites (10–14). The efficiency of malaria transmission-blocking and the stability of association with the vector are important for potential application for malaria control. Notably, endosymbiotic microbes that have high-fidelity vertical transmission have an inherent ability to maintain their prevalence across mosquito generations and are therefore likely to be sustainable and cost-effective as malaria transmission blockers. Recently a native endosymbiont, *Microsporidia MB*, was discovered in *Anopheles* mosquitoes and shown to impair the transmission of the *Plasmodium falciparum* parasite by *Anopheles arabiensis* mosquito (14). Unlike many previously reported microsporidians which are pathogens of mosquitoes (15), *Microsporidia MB* does not have any apparent negative fitness effects on its host and is vertically transmitted from mother to offspring (14). *Microsporidia MB* is also sexually transmitted between mating pairs (16). Together, these characteristics could enable its spread through mosquito populations (14). Recently, we have shown that *Microsporidia MB* is predominantly localized in the reproductive organs which is relevant for vertical transmission from mother to offspring and horizontal sexual transmission (16–18). However, the precise mechanisms involved in the vertical transmission of *Microsporidia MB* have not been elucidated. Understanding the strategies employed by this symbiont to vertically infect *Anopheles* mosquitoes will be important to determine the factors limiting infection rates and specificity to different malaria vector species and populations. Understanding these factors could facilitate strategies to sustainably increase *Microsporidia MB* infections in natural populations to significantly reduce malaria infections.

In this study, we investigated how the microsporidian symbiont is transferred from mother to offspring by studying its localization within the cells of the *Anopheles* mosquito ovary. We report that *Microsporidia MB* is present in the germarium and infects the germline stem cells from where it enters the cystocytes as a result of stem cell division. As the cystocytes develop into follicles, *Microsporidia MB* infection intensities increase first in the nurse cells and then in oocytes through active symbiont cell division. The average rate of transmission from stem cells to follicles (>96%) is higher than the average rate of observed adult to adult vertical transmission, suggesting that some *Anopheles* mosquitoes lose their symbiont infections over development.

## Results

### *Microsporidia MB* is vertically transmitted in the *An. arabiensis* germarium

*Microsporidia MB* was localized in *An. arabiensis* female ovaries using fluorescence in situ hybridization (FISH) microscopy with a *Microspordia MB* specific probe (14). *An. arabiensis* ovaries comprise tens to hundreds of polytrophic ovarioles with each ovariole composed of a germarium, the secondary, and the primary follicles. The germarium is located at the distal tip of the ovariole and contains 2-4 germline stem cells (GSCs). As each GSC divides, it produces an identical GSC and differentiating germline cell called a cystoblast. As a cystoblasts progress through the germarium, it undergoes mitotic divisions in combination with incomplete cytokinesis resulting in a follicle that has seven nurse cells connected through cytoplasmic bridges and one oocyte (Fig 1A&B). Through the detection of newly synthesized DNA, we show evidence of newly formed cystoblasts following stem cell divisions in the germarium of emerged two-day-old mosquitoes (Fig 1C). We demonstrate the localization of *Microsporidia MB* in the germarium, inside the germline stem cells from where it is transferred to cystocytes during cell division. We also found *Microsporidia MB* infections in the secondary and primary follicles (Fig 1D-F). In the primary follicles, the symbiont was present in the developing oocytes and the nurse cells in high abundance but not in the follicular epithelial cells (Fig. 1D-F).

**Figure 1.**
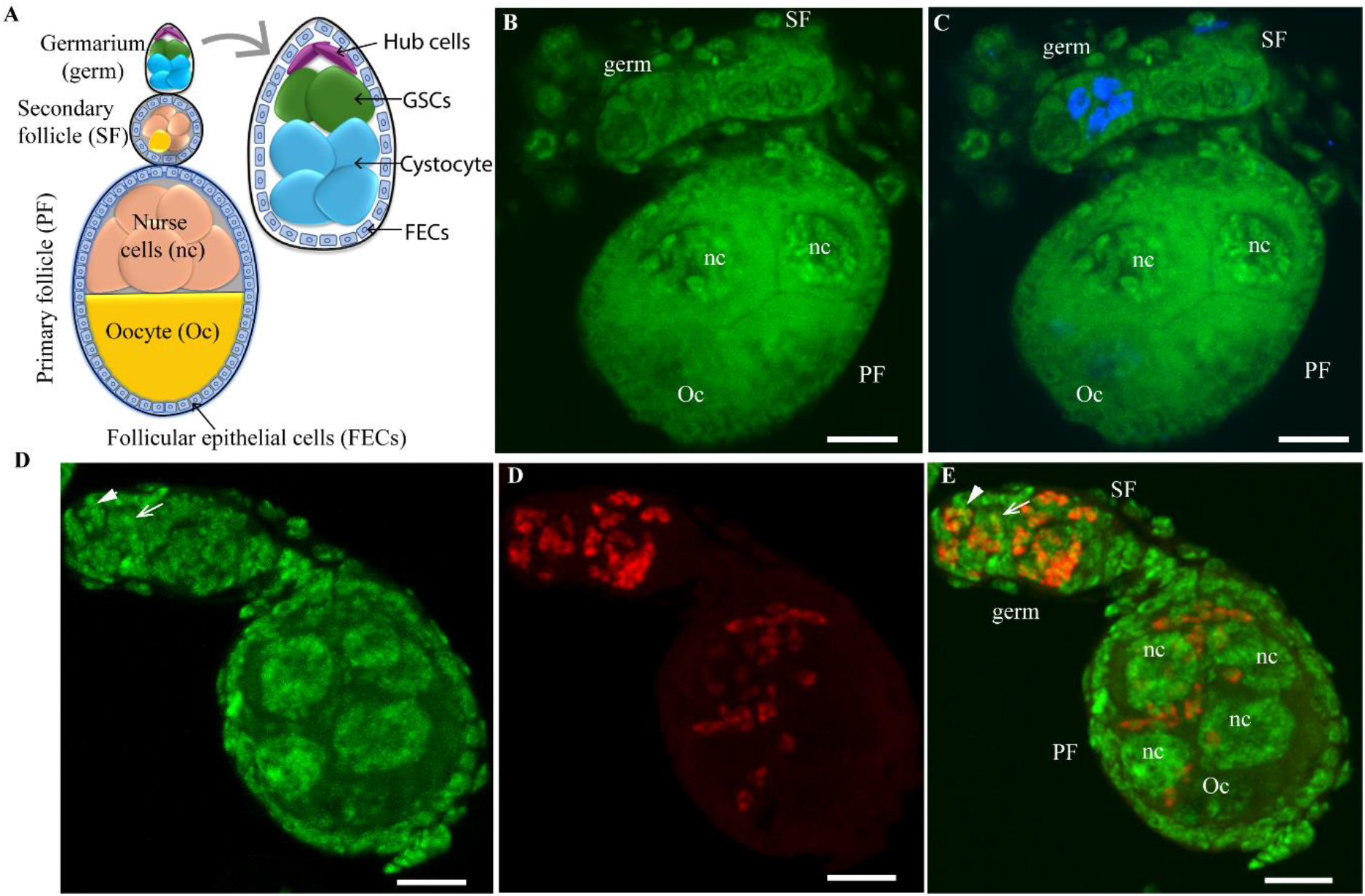
*Microsporidia MB* localization in the polytrophic *An. arabiensis* ovariole. (A) Schematic representation of the ovariole structure showing the germarium, the secondary follicle, the primary follicle with the oocyte and nurse cells in one chamber, and the epithelial follicular cells. The germline stem cells (GSCs) and cystocytes are located in the germarium. (B) *Microsporidia MB* negative control ovariole of a two-day-old *An. arabiensis* mosquito showing the localization of stem cell daughter cells (in blue) following cell divisions within the germarium, (C). (D-F) *Microsporidia MB*-infected ovarioles of *An. arabiensis* mosquito (Green represents Sytox Green DNA staining, red represents *Microsporidia MB-*specific FISH probe, and a merged image of the green and red channel). The symbiont is present in the germarium (germ) in the GSCs (white arrowheads pointing the GSC nucleus) and cystocytes (white arrow pointing the cystocyte nucleus), the secondary follicle (SF), and the primary follicle (PF) i.e., the nurse cells (nc) and the oocyte (Oc). Scale = 10µm. (Images are representative pictures of >100 independent observations).

### *Microsporidia MB* dynamics during egg development and maturation

Differentiation of the primary follicle into a mature egg within the ovariole in Anopheline mosquitoes is characterized by seven stages; stages I-III (previtellogenesis) and stages IV-VII (vitellogenesis). The vitellogenic stages IV-VII only occur after the female mosquito takes a blood meal. To determine *Microsporidia MB* growth dynamics in the ovariole during development, we performed FISH microscopy for ovarioles at different stages of development and quantified relative *Microsporidia MB* intensities in their oocytes. Relative quantification was achieved by measuring the area covered by *Microsporidia MB* fluorescence signal in confocal microscopy images in relation to the entire area covered by the oocyte. We analyzed *Microsporidia MB* localization patterns in the germarium, the secondary follicle, and the primary follicle across all ovariole developmental stages (Figure 2A). In the early previtellogenic stages (stages I and II), the symbiont is evenly distributed in the nurse cells and oocyte of the primary follicle (Figure 2A). In the late previtellogenic stage and vitellogenic (stages III-VI), however, *Microsporidia MB* accumulates in the developing oocyte while decreasing in the nurse cells (Figure 2A). Once the egg reaches maturity and takes a typical cigar shape following the degeneration of nurse cells, an even distribution of *Microsporidia MB* cells throughout the embryo is observed (Figure 2A, stage VII).

**Figure 2.**
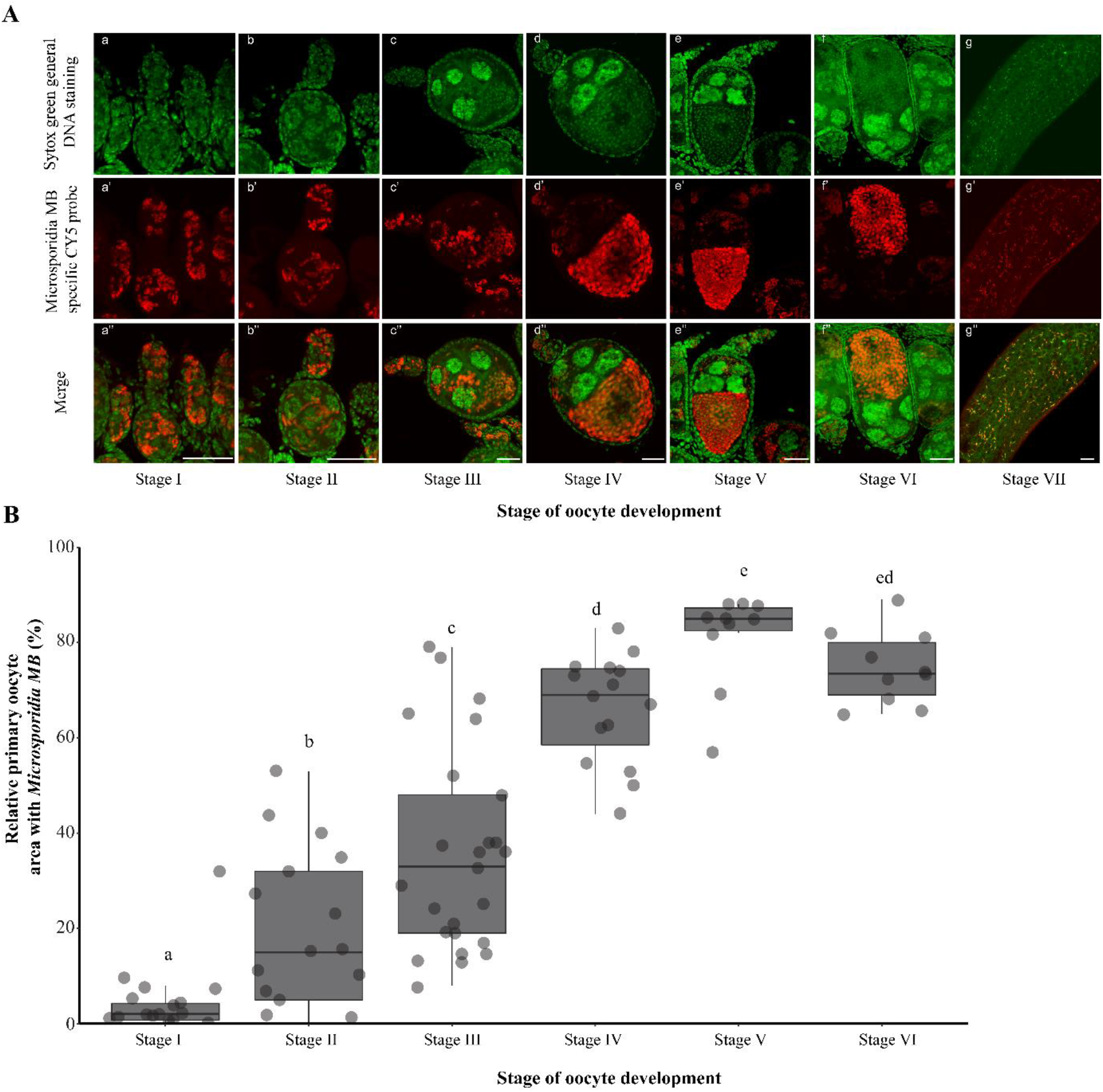
Localization of *Microsporidia MB* within the germarium and follicles across the different developmental stages of *An. arabiensis* ovariole. (A) *Microsporidia MB* is spread throughout the primary follicle during the previtellogenic phase of development (stages I-III). As development progresses, microsporidia symbiont cells accumulate in the anterior end of the primary follicle oocyte in high densities during the vitellogenic phase (stages III-VI). The upper panels (a-g) represent Sytox Green DNA staining signal, the second row of panels (a’-g’) represents *Microsporidia MB-specific* CY-5 staining signal and the panels in the third row (a’’-g’’) represent merged images of both signals. Scale = 25µm. (B) The relative area of the primary follicle oocyte occupied by *Microsporidia MB* symbiont. *Microsporidia MB* occupies a small area of the primary follicle oocyte in the previtellogenic stages of development and increases significantly during the vitellogenic stages of development (χ^2^(5) = 75.04, p < 0.01).

Within the primary follicle, *Microsporidia MB* was mainly localized in the oocyte of the maturing egg. The intensity of *Microsporidia MB* infections in previtellogenic stage oocytes is low since the average relative area of the primary oocyte covered by *Microsporidia MB* is less than 10% (Figure 2B). However, the infection intensities significantly increase during vitellogenic stages of development to occupy a relative area of more than 80% of the entire primary follicle oocyte in stage V of egg development (χ^2^(5) = 75.25, p < 0.01). This increase of *Microsporidia MB* intensities inside the primary follicle oocyte suggests active proliferation of the symbiont or its deposition from the nurse cells.

### *Microsporidia MB* actively divides in the oocyte

We observed an increase of *Microsporidia MB* intensities in oocytes of the primary follicles across the vitellogenic developmental stages (Fig. 2). To establish how *Microsporidia MB* intensities increase in the developing primary follicles, we investigated the physiological state of *Microsporidia MB* cells within the nurse cells and oocyte in the primary follicles and stained for actively dividing cells in the ovariole. We identified different sizes of *Microsporidia MB* cells inside the primary follicles ranging from smaller cells with a size of about 2μm to larger cells with a size of about 5μm (Fig 3A) suggesting that the symbiont exists in multiple forms in the follicle. We also observed *Microsporidia MB* cells in the process of fission (merogony) in the oocyte, which suggested that the symbiont was actively dividing within the developing oocyte (Fig 3B). To confirm occurrence of *Microsporidia MB* proliferation in the ovariole, we stained for actively dividing cells in *Microsporidia MB-*infected ovarioles using Click-iT EdU cell proliferation assay that detects newly synthesized DNA. We detected new *Microsporidia MB* cells mostly in the oocyte of the primary follicle but also in the secondary follicle and the germarium (Figure 3C). The size of the newly synthesized *Microsporidia MB* nuclei was consistent with the staining we observed with Sytox Green DNA staining.

**Figure 3.**
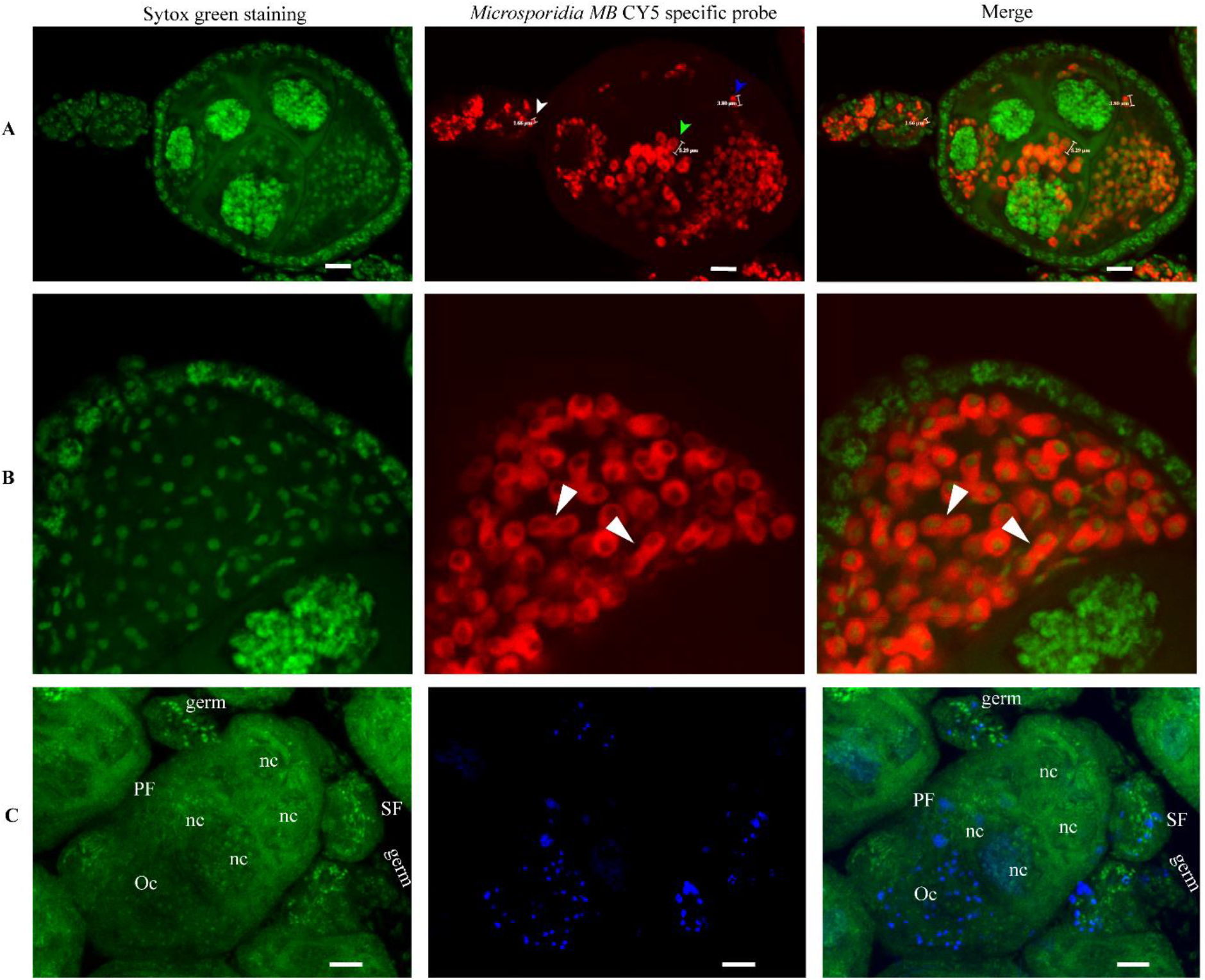
*Microsporidia MB* development in the ovariole. (A) Different sizes of *Microsporidia MB* cells exist in the ovariole ranging from 2μm in the germarium and secondary follicle (white arrowhead) to 5μm in the nurse cells and oocyte of primary follicle, green and blue arrowheads, respectively. (B-C) Evidence of *Microsporidia MB* proliferation in the ovarioles. *Microsporidia MB* cells in the process of fission in the oocyte (B, white arrowheads) and the detection of newly synthesized DNA in actively proliferating cells (C). Green represents the general DNA staining signal (Sytox Green), red represents *Microsporidia MB* staining signal (*Microsporidia MB-specific* CY-5 probe), and blue shows the signal of newly synthesized DNA (Click-iT Edu staining).

### Vertical transmission rate of *Microsporidia MB* into primary follicles

We quantified *Microsporidia MB* infection rates of primary follicles in the ovaries to establish the vertical transmission rate of *Microsporidia MB* into developing eggs in infected ovaries. We found that a small number of primary follicles (3%) were not infected with *Microsporidia MB* while a majority (97%) of the primary follicles in infected ovaries had *Microsporidia MB* (Fig 4A). Interestingly, in ovarioles whose primary follicles were not infected with *Microsporidia MB* (3%), the symbiont was consistently present in the germarium and the secondary follicles (Fig. 4B) suggesting that early stem cell divisions could occasionally not transfer *Microsporidia MB* to the follicles. To establish whether this phenomenon could explain the imperfect vertical transmission of *Microsporidia MB*, we compared the primary follicle infection rate and *Microsporidia MB* prevalence in emerging adult F1 offspring of *Microsporidia MB*-infected isofemale mothers. We found that whereas 97% of the primary follicles were infected with *Microsporidia MB*, only 61% of emerging adult F1 offspring were infected with *Microsporidia MB* (Fig. 4A), suggesting that *Microsporidia MB* can be lost during pre-imaginal stages of development. In all *Microsporidia MB-*infected *An. arabiensis* female gonads we investigated, we found the symbiont to be localized in both ovaries indicating even distribution of the symbiont in infected gonads (Figure 4C).

**Figure 4.**
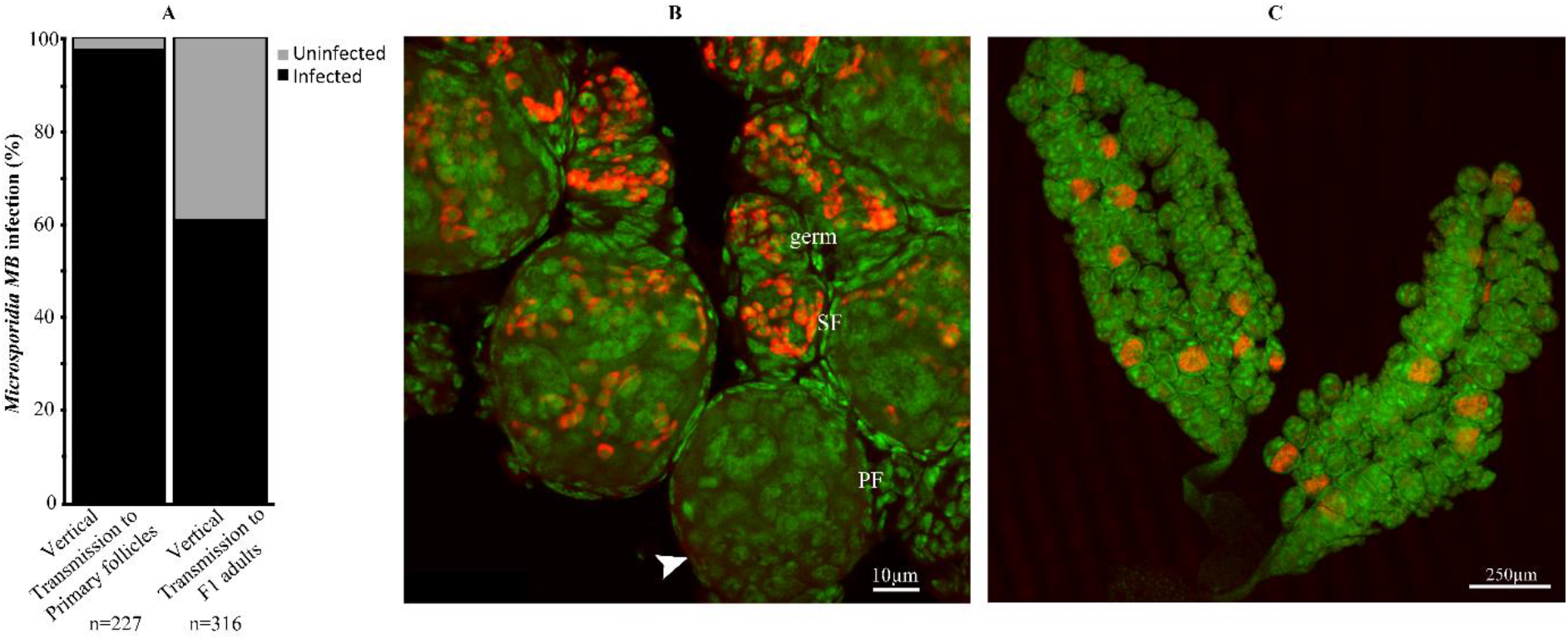
*Microsporidia MB* infection rates in primary follicles and vertical transmission rates. *Microsporidia MB* distribution in infected female ovaries was investigated and compared to vertical transmission rates. (A-B) A majority (97%) of the primary follicles within infected ovarioles had *Microsporidia MB* while the symbiont was completely absent in 3% of the primary follicles (B, white arrowhead). However, the germarium (germ) and secondary follicles (SF) of the 3% primary follicles that lacked *Microsporidia MB* were consistently infected with *Microsporidia MB*. *Microsporidia MB* was detected in 61% of the adult offspring of *Microsporidia MB-*infected mothers. In all infected female gonads investigated, the symbiont is present in both ovaries (C). Green represents general DNA staining (Sytox Green) while red represents *Microsporidia MB-specific* CY-5 staining.

## Discussion

Successful deployment of a novel *Microsporidia MB*-based approach for controlling malaria will depend on the ability of the microsporidian symbiont to invade, spread, and be maintained in target *Anopheles arabiensis* vector populations. While *Microsporidia MB* has high rates of maternal transmission in *An. arabiensis*, these rates are variable. The rate of maternal transmission is critical for determining the prevalence of *Microsporidia MB* in a population of *Anopheles* mosquitoes and the impact on *Plasmodium* (19). In this study, we established the mechanistic basis of *Microsporidia MB* vertical transmission, which occurs through the acquisition of *Microsporidia MB* by *Anopheles* cystocyte progenitors from the germline stem cells following cell division. As the daughter stem cell cystocytes differentiate into follicles, *Microsporidia MB* establishes itself in the nurse cells and ultimately the oocyte and egg. Our findings suggest that *Microsporidia MB* becomes established in the *Anopheles* germline early during development and this seems to be highly efficient, as only 3% of primary follicles do not aquire a *Microsporidia MB* infections. We didn’t find any evidence of the germline becoming re-infected from infections of somatic cells, but we do not rule out that this could be occurring. In particular, we have previously observed that *Microsporidia MB* infections acquired via sexual transmission can subsequently be vertically transmitted. Vertical transmission of microsporidian symbionts through egg infections has been reported in different species of mosquitoes and in the amphipod *Gammarus duebeni* (20–24). However, symbiont infections in these hosts were only observed in the oocyte and epithelial follicular cells but not in the germline stem cells suggesting that the mechanism of vertical transmission is through somatic cell infections and not germline stem cell infections as we found for *Microsporidia MB*. Somatic cell infections and subsequent transovarial transmission has been demonstrated for *Edhazardia aedis* in *Aedes aegypti* and *Amblyospora* sp. in *Aedes cantator*, *Aedes triseriatus* and *Culex salinarius* when horizontally (orally) acquired spores infect oenocytes and spread to the ovaries where they proliferate and form transovarial spores (15,20,22,25). Similarly, vertical transmission of a microsporidian symbiont of *Gammarus duebeni* occurs when the symbiont invades and proliferates in follicular cells before spreading into the developing oocytes for transmission to offspring (21).

The mechanistic basis of *Microsporidia MB* transmission in *Anopheles arabiensis* is strikingly similar to *Wolbachia* infections in other insects including *D. melanogaster* and *Aedes* mosquito species where the endosymbiont is in the germ line stem cells and is vertically transferred to the progeny following stem cell divisions (12,26–28). For *Wolbachia*, there is apparently an additional vertical transmission route, which involves infection of the follicle from the surrounding somatic cells. It is possible that this additional infection route is important for maintaining near perfect rates of vertical transmission, which are required for an endosymbiont that is entirely dependent on vertical transmission (28,29). In this study, *Microsporidia MB* was not observed in the somatically derived epithelial follicular cells that surround the egg chamber. *Wolbachia* infections of the germline stem cells of *Drosophila* are capable of influencing mitotic processes to promote more egg production (30,31). Although *Microsporidia MB* does not apparently affect egg production in *Anopheles arabiensis* (Herren et al., 2020), its high infection intensity in the germline stem cells and developing oocytes may have consequences for cellular processes and embryonic development. It is notable that the mechanistic basis of *Microsporidia MB* vertical transmission is very different from another insect endosymbiont, *Spiroplasma*. *Spiroplasma* is absent from the germline stem cells but gains access to the vitellogenic oocyte during vitellogenesis by co-opting the yolk transport system (32). The existence of *Microsporidia MB* in the stem cells and oocyte of developing eggs in *An. arabiensis* provides strong evidence for a highly specialized vertical transmission strategy for *Microsporidia MB* in *An. arabiensis*. This high level of specialization may explain why *Microsporidia MB* has so far only been found in *Anopheles* mosquitoes (16).

Similar to other microsporidians, *Microsporidia MB* is a unicellular obligate intracellular organism that can only replicate and survive inside the host cells and only exist as spores outside the host cells (33). Our findings demonstrate that cystocytes, nurse cells, and oocytes are among the suitable host cell types for this symbiont within the *An. Arabiensis* host. We identified different sizes of *Microsporidia MB* cells in the ovariole suggesting that *Microsporidia MB* is transitioning through different life cycle stages in this tissue. Further, we found evidence that the symbiont actively proliferates inside the primary follicle oocyte through the detection of *Microsporidia MB* cells undergoing merogony and the presence of newly synthesized *Microsporidia MB* DNA. This proliferation was observed after mosquitoes had a successful blood meal suggesting that *Microsporidia MB* proliferation is coordinated with host cellular processes. Similarly, sporulation of *Amblyospora* sp. in *Culex salinarius* and *Culiseta incidens* only occurs after the mosquitoes take a blood meal and it coincides with the maturation of the oocytes (20,34). The development of the microsporidian symbiont of *Gammarus duebeni* is also highly coordinated with the host reproductive cycle (21). The controlled proliferation of symbionts within insect hosts is important in mitigating any fitness costs to the host (21,35,36). Unlike other pathogenic microsporidian symbionts that kill their hosts (33), *Microsporidia MB* has no apparent fitness costs to the host despite attaining high infection intensities in the oocyte of the developing egg (14). For vertically transmitted symbionts, there’s a strong selection for avirulence to the host because the symbiont’s success is associated with the host’s reproductive success (37,38). Therefore, the specialized vertical transmission of *Microsporidia MB* through infection of oocytes of developing eggs of *An. arabiensis* suggests that avirulence has been selected for during evolution leading to its low pathogenicity to the host.

The presence of the microsporidian symbiont in most (97%) of the developing eggs implies a potential for high vertical transmission rates. However, in our experiments, we observed an average of 61% vertical transmission rate from infected mothers to adult F1s, which was lower than the infection rate in eggs that were developing in the ovary. Previously, we reported marginally higher transmission rates of *Microsporidia MB* from infected mothers to their offspring (14). The absence of the symbiont in some of the primary follicles only partially explains the imperfect vertical transmission rate. Additionally, other factors such as the host immune responses and environmental factors may be responsible for the loss of infections during mosquito development. As demonstrated in other insect-symbiont associations (39–41), the presence of *Microsporidia MB* may elicit *A. arabiensis’* immune response which then directly interferes with the establishment of the symbiont before it colonizes the primary sites of infection where it may be protected from the host immune responses. Also, environmental factors such as temperature and humidity that are known to influence the occurrence and transmission of different endosymbionts including microsporidian symbionts in different insects are likely to affect the establishment of *Microsporidia MB* (42–44).

Our study has shown a specialized vertical transmission route of *Microsporidia MB* in *An. arabiensis* that is relevant in developing a symbiont-based strategy to control malaria. These findings will serve as a basis for investigations on different *Microsporidia MB* strains and transmission phenotypes in other Anopheline vectors aimed at maximizing *Microsporidia MB* vertical transmission rates for the development of a viable strategy for symbiont-based malaria transmission blocking.

## Methods

### Mosquito collection and rearing

Gravid *An. gambiae* s.l mosquitoes were collected from the Ahero irrigation scheme in Kenya in 2022 and 2023 and transported to icipe’s Duduville campus in Nairobi Kenya. The mosquitoes were then maintained in the insectary at 27 ± 2.5°C, humidity 60–80%, 12-h day and 12-h night cycles and induced to oviposit through the provision of oviposition cups with a wet cotton wool lined with a filter paper. After ovipositing, DNA was extracted from the females and used for species identification and screening for *Microsporidia MB* infections using PCR (14). Offspring from *Microsporidia MB-*infected and uninfected female *An. arabiensis* mosquitoes were then reared in the insectary as described above for experimental investigations. Blood feeding of mosquitoes to initiate the processes of vitellogenesis and oogenesis was conducted as previously described (Makhulu et al., 2024).

### FISH localization of *Microsporidia MB* in *An. arabiensis* ovaries

Fluorescent in-situ hybridization was conducted to localize *Microsporidia MB* in the ovarioles of newly emerged 2-day-old and 2-5 days post-blood-fed *An. arabiensis* mosquitoes. Ovaries of *Microsporidia MB*-infected and uninfected (controls) were dissected in 1x PBS and fixed overnight in 4% Paraformaldehyde (PFA) at 4 °C then rehydrated in 50% ethanol for 30 minutes, followed by a washing step in 1x PBS for 30 minutes. FISH was conducted as previously described to localize *Microsporidia MB* within the ovarioles (14,18). Briefly, hybridization was done by incubating the tissues at 50°C overnight in 100μl of hybridization mix (hybridization buffer i.e., dH2O, 5M NaCl, 1M Tris/HCl [pH=8], and 10% SDS, 0.5μM final concentration of the *Microsporidia MB* specific CY5 probe (10), and SYTOX Green general DNA staining. After staining, the samples were washed twice with 100μl of wash buffer prewarmed at 50°C (dH2O, 5M NaCl, 1M Tris/HCl [pH=8], 0.5M EDTA, and 10% SDS). The tissues were subsequently mounted on glass slides using fluoromount mounting media and visualized immediately using a Leica SP5 confocal microscope (Leica Microsystems, USA). The images were analyzed with the ImageJ 1.50i software package (31).

### Click-iT® EdU staining of actively dividing cells in *An. arabiensis* germarium

Ovaries of *An. arabiensis* females were dissected in 10μM EdU in 1x PBS solution and labeled in this solution for one hour, then fixed in a 3.7% PFA solution for 30 minutes. Subsequently, they were washed twice with 500µL of 3% BSA in 1x PBS and permeabilized using 0.5% Triton® X-100 in 1x PBS for 20 minutes at room temperature before washing twice with 500μL of 3% BSA in 1x PBS solution. The ovaries were then stained and protected from light for 2 hours in freshly prepared 500μL of Click-iT® Plus reaction cocktail prepared as per the manufacturer’s specifications. Once the Click-iT® staining was complete, the tissues were washed in with 500μL BSA in 1x PBS solution and counterstained with SytoGreen general DNA stain (5 µg/ml) before mounting on a slide using fluoromount mounting media and imaging on the Leica SP5 confocal microscope (Leica Microsystems, USA). The images were analyzed with the ImageJ 1.50i software package (31).

### Quantification of *Microsporidia MB* in *An. arabiensis* Ovaries

ImageJ software was used to independently measure the area occupied by *Microsporidia MB* in the primary follicle oocyte by measuring fluorescence intensity. The images were first converted into an 8-bit grayscale binary scale. The regions of interest (ROI) around each of these structures were defined using the follicular epithelial cells so that only the pixels within the area of interest were selected and included in the final measurements. Subsequently, we measured the proportion of the area that was occupied by *Microsporidia MB* cells within each of the ROIs as previously described (45). The relative relative oocyte area covered by *Microsporidia MB* across the different stages of egg development was compared using Kruskal-Wallis H test. R software was used for statistical analysis.

### *Microsporidia MB* infection rates in the ovaries

To determine *Microsporidia MB* infection rates in the ovaries, a complete set of eight ovaries was obtained from select offspring of wild-caught *Microsporidia MB*-infected mothers that were collected from different locations within Ahero. The ovaries were stained by FISH as described above and the presence or absence of *Microsporidia MB* in both ovaries was recorded. *Microsporidia MB* infection rates in the primary follicles were determined by randomly selecting and counting the number of infected and uninfected follicles in the eight pairs of ovaries.

### Vertical transmission rate of *Microsporidia MB*

Field-collected gravid females were induced to oviposit as described above and then screened for *Microsporidia MB* infections using PCR (14). Eggs from *Microsporidia MB-*infected mothers were hatched in the insectary and maintained at a temperature of 27 degrees and fed on tetramin fish food until they pupated. The isofemale line pupae were subsequently transferred into cages where they emerged and maintained on a sugar diet. DNA was extracted from these offspring two days after emergence and used to screen for *Microsporidia MB* with PCR (14,18). The number of infected adult offspring per isofemale line was expressed as a percentage.

## Acknowledgment

We acknowledge the support provided by the *icipe* insectary team (Jeniffer Thiong’o, Peris Wambui), fieldwork facilitation by Gerald Ronoh, and project administration by Faith Kyengo.

## Data accessibility

The relative area of the primary oocyte infected with *Microsporidia MB* across developmental stages of the ovariole, the primary follicle infection rate, and vertical transmission rate of *Microsporidia MB* to adults data are available in the Figshare Digital Repository at https://doi.org/10.6084/m9.figshare.25846351

## Author contributions

TOO – Conceptualization, investigation, and data collection, data curation, validation, visualization, methodology, formal analysis, writing – original draft, writing – review & editing.

EMM, PCR, SM, FGO, JG – Investigation and data collection, Data curation, validation, visualization, methodology, writing – review & editing.

JKH – Conceptualization, data curation, formal analysis, funding acquisition, methodology, supervision, validation, visualization, writing – original draft, writing – review & editing.

